# SpatialFinder: A Human-in-the-Loop Vision-Language Framework for Prioritizing High-Value Regions in Spatial Transcriptomics

**DOI:** 10.1101/2025.08.16.670684

**Authors:** Jonathan Xu, Michelle Jiang, Shunsuke Koga, Nancy Zhang, Zhi Huang

## Abstract

Sequencing an entire spatial transcriptomics slide can cost thousands of dollars per assay, making routine use impractical. Focusing on smaller regions of interest (ROIs) based on adjacent routine H&E slides offers a practical alternative, but there is (i) no reliable way to identify the most informative areas from standard H&E images alone; and (ii) limited solutions for clinicians to prioritize the microenvironment of their own interests. Here we introduce **SpatialFinder**, a framework that combines a biomedical vision-language model (VLM) with a human-in-the-loop optimization pipeline to predict gene expression heterogeneity and rank high-value ROIs across routine H&E tissue slides. Evaluated across four Visium HD tissue types, SpatialFinder consistently outperforms baseline VLMs in selecting regions with high cellular diversity and tumor presence, achieving up to 89% correlation with ground truth rankings. These results demonstrate the potential of human-AI collaboration to make spatial transcriptomics more cost-effective and clinically actionable.

## 1. Introduction

Spatial transcriptomics (ST) enables *in situ* quantification of gene expression across intact tissue sections, preserving spatial context to map transcriptional activity within anatomical niches^1^. This spatial resolution makes it possible to distinguish expression patterns in specific regions, such as a tumor’s core versus its invasive edge, revealing interactions that bulk or single-cell RNA-seq would miss^2^. Modern ST technologies enable gene expression profiling across entire tissue sections^3^, but practical adoption is hindered by high cost and scalability challenges^4^. Sequencing a single slide can cost up to $15,000 using advanced platforms like 10xGenomics Visium HD^5,6^. These assays remain too expensive and thus impractical for usage in widespread research or clinical settings^7,8^. Moreover, large portions of a tissue slide may offer little informative value or repeated patterns, leading to significant investment in sequencing regions that contribute minimal biological insight^9,10^. This inefficiency motivates the need for methods to intelligently *target* sequencing to the most informative subregions of a tissue sample^11^. By reliably identifying smaller areas that reflect the tissue’s gene expression diversity, we could focus sequencing efforts where they matter most, drastically reducing costs while preserving essential molecular information^12^. Such an approach would make spatial transcriptomics much more accessible for both research and clinical use in the future.

In the past decade, artificial intelligence (AI) has made rapid strides in analyzing whole-slide histology images^13^. At the forefront of this progress are vision-language models (VLMs), which learn joint representations of images and text^14^. In pathology, VLMs can localize regions matching natural language prompts, such as “tumor region with dense lymphocyte infiltration”, even without explicit training on those patterns^15,16^. By aligning semantic concepts with visual features, VLMs offer a flexible way to detect novel or rare tissue patterns that may reflect underlying molecular or clinical features^17,18^.

Recent efforts in both bioinformatics and AI have explored strategies to reduce the experimental burden of spatial transcriptomics^19^. Image-to-omics prediction models have been developed to computationally infer spatial gene expression patterns directly from histology images^20^. For example, convolutional neural networks and graph neural networks have been applied to predict transcriptomic profiles in unseen tissue regions, offering a promising *in silico* alternative to exhaustive sequencing^21^. However, a limitation of purely computational approaches is the uncertainty in predictions, which still benefits from validation via actual sequencing in critical regions^22^. Other related work has focused on identifying regions-of-interest in pathology slides using weakly supervised learning and anomaly detection, to guide pathologists or downstream analyses^23,24^. While these data-driven AI models are increasingly powerful, they often act as black boxes, lacking transparency and sometimes producing clinically implausible results^25,26^. Most importantly, expert clinicians have limited access to steer or evolve medical AI systems in line with clinical and research needs^27^. To keep AI tools accurate and trustworthy in medicine, especially in the scenario of choosing ROI for reducing ST costs, a human-in-the-loop component is essential: integrating expert feedback throughout training and deployment can align AI models’ predictions with clinicians’ expectations^28^. To let clinicians be part of this process, active learning is one effective strategy, where the model selectively queries experts to label uncertain or informative samples and retrains accordingly^29^. This process focuses the model on clinically relevant features with far fewer annotations than traditional methods^30^. Studies have shown that such human-guided pipelines not only improve performance but also reduce labeling effort and training time^31,32^.

Unlike previous approaches, our proposed SpatialFinder identifies optimal ROIs for spatial transcriptomics by leveraging vision-language features to capture semantic heterogeneity. Uniquely, it incorporates a human feedback loop to iteratively refine predictions. Furthermore, by benchmarking selected subregions against ground truth spatial omics data, we introduce an evaluation framework for region selection, a gap not explicitly addressed in existing studies. In evaluations across four Visium HD tissue types, SpatialFinder demonstrated up to 89% agreement with ground truth region rankings and achieved substantially greater spatial coverage of top regions compared to leading baselines. Notably, just 5–10 minutes of expert feedback yields up to a 50 percentage-point improvement in overlap with ground truth among the highest-ranked regions, underscoring the practical impact of human-in-the-loop refinement.

## 2. Methods

We present a novel framework that combines a vision-language model with human-in-the-loop supervision to identify informative subregions in histology images for spatial transcriptomics. To our knowledge, this is the first approach to integrate a foundation VLM with expert pathologist input for region selection in this context. The method consists of two parallel components: (i) a pathologist-guided cellular classifier trained on hematoxylin and eosin (H&E) stained slides for fine-grained, single-cell recognition of histological patterns, and (ii) a VLM that encodes image patches into high-dimensional embeddings for zero-shot interpretation of complex tissue structures. By combining expert-labeled cellular features with the semantic generalization of a VLM, our system robustly identifies regions likely to capture diverse and relevant gene expression profiles.

### 2.1. Data Curation and Preprocessing

We curated a dataset of high-resolution Visium HD H&E-stained whole-slide images (WSIs) from 10xGenomics, spanning four tissue types: prostate cancer^33^, lung cancer^34^, colon cancer^35^, and non-cancerous kidney^36^. Each slide includes a defined capture area with ground truth spatial transcriptomics data. We first perform automated cropping to isolate the tissue capture region from the full WSI, focusing our analysis on the area with corresponding gene expression data.

After cropping, we perform comprehensive cell nucleus segmentation on each tissue region. We apply a pre-trained StarDist deep learning model to detect and delineate each cell nucleus in the image^37^. This yields a detailed segmentation mask marking the precise location and boundaries of each nucleus in the tissue. The resulting mask serves as the basis for subsequent pathologist-guided cell type annotation and classification.

### 2.2. Morphological Feature Extraction with VLM

To capture high-level morphological features, we use a state-of-the-art biomedical vision-language model. Each cropped tissue image is divided into a grid of small patches (224×224 pixels each), with an initial saturation-based threshold excluding empty background regions for efficiency. The remaining patches are processed with the Multimodal transformer with Unified maSKed modeling (MUSK), a transformer-based VLM pre-trained on large-scale pathology image-text data^38^. MUSK generates a 1024-dimensional embedding per patch, encoding rich morphological semantics for zero-shot inference. To ensure feature quality, we apply a two-stage filter: first removing background patches (high mean intensity, low variance), then excluding any remaining low-variance tissue.

### 2.3. Pathologist Classification via Active Learning

To achieve fine-grained cellular classification, we employ a hybrid active learning workflow powered by NuClass, a custom classification tool integrated within our in-house TissueLab software platform. This workflow begins with an automated, zero-shot labeling of all segmented nuclei. We use the text encoder from a fine-tuned Pathology Language-Image Pretraining (PLIP) model^39^ to embed pathologist-defined text labels (e.g. “tumor cell,” “immune cell”) as feature vectors. Each nucleus is also embedded via the PLIP model and assigned the label with the highest cosine similarity, enabling rapid, semantic classification without manual annotation.

This initial, AI-generated map serves as a starting point for an expert pathologist, who uses TissueLab’s interface to label small, representative subsets of nuclei. These expert annotations, comprising each cell’s image embedding and its correct label, form a training set for a supervised XGBoost classifier^40^. Once trained on this curated dataset, the model is reapplied to predict labels for all nuclei across the tissue. The pathologist can then iteratively refine the results by providing more labels, particularly in areas of uncertainty. With each cycle, the XGBoost classifier is retrained on the cumulative set of annotations, rapidly improving its accuracy and converging into a robust model with only a few hundred expert labels. This process produces a spatially-resolved cell type map for hundreds of thousands of cells, effectively blending large-scaled automated analysis with precise expert knowledge.

### 2.4. A Hybrid Model for Identifying Regions of Interest

To prioritize ROIs for sequencing, we use a hybrid model that integrates VLM-based visual features with the pathologist-tuned cell type map. We scan the tissue using an 8 × 8 patch sliding window (∼ 500 × 500μm, or 0.5% of the 6.5 × 6.5mm Visium HD total area), generating approximately 10,000 ROIs per slide. For each resulting ROI, we compute two distinct scores: a Regional Diversity Score for unsupervised discovery of biological complexity, and a Targeted Conditional Score for hypothesis-driven searches.

The **Regional Diversity Score** measures biological heterogeneity using: (i) visual diversity, and (ii) cellular diversity. For visual diversity, patch embeddings (1024-D) are reduced to 30-D using UMAP^41^, and median pairwise Euclidean distance is computed. Higher values indicate morphologic variation (e.g., mixed glandular, stromal, and tumor areas). For cellular diversity, we compute Shannon entropy^42^ of predicted cell types for the nuclei within each ROI to reflect its compositional richness. The final score is *S*_Diversity_ = (1−*w*)*·*Score_cellular_+*w·*Score_visual_.

The **Targeted Conditional Score** identifies ROIs matching a user-defined biological query, combining (i) text-guided VLM similarity and (ii) target cell abundance from the classifier. For the former, a prompt is embedded and compared (via cosine similarity) to each patch in the ROI. We use the 90th percentile of these scores, scaled exponentially to emphasize strong matches. For the latter, we derive the log-transformed count of the relevant cell type (e.g., tumor) based on classifier outputs. The final targeted score is similarly computed as *S*_Targeted_ = (1 − *w*) *·* Score_count_ + *w ·* Score_vlm_, where *w* is a tunable weight.

Each metric is used independently to rank all ROIs, and top-ranked regions can be selected for downstream sequencing. The top selections for each tissue slide are highlighted below:

### 2.5. Experimental Validation

To validate our method, we benchmarked its ROI rankings against ground truth derived from the Visium HD feature matrix, which provides gene expression data in a grid of 8 × 8μm bins, or “spots”, each approximately the size of a few cells^a^.

We defined cell type-specific marker genes based on canonical sets curated from biological literature and prior databases^43–50^,^b^. Marker selection was tailored to each tissue type:

- **Lung and colon cancer**: Tumor cells were identified using epithelial and carcinoma markers such as *EPCAM* and *CEACAM5*. Immune cell types were defined by *PT-PRC* (pan-immune), *CD3D* and *CD8A* (T cells), and *MS4A1* (B cells). Stromal and endothelial populations were marked by *COL1A1* and *PECAM1*, respectively.
- **Prostate cancer**: Tumor cells were labeled using *AMACR* and *KLK3* (encoding *PSA*). Basal and luminal epithelial cells were identified with cytokeratins like *KRT5* and *KRT8*.
- **Non-cancerous kidney tissue**: Cell types were annotated by functional nephron markers—*NPHS1* (podocytes), *LRP2* (proximal tubule), and *UMOD* (loop of Henle).

We used the 8μm ST spots as the basic unit of ground truth. Each spot was classified by computing the mean expression of each marker gene set and assigning it to the cell type with the highest score above a minimum threshold. This yielded a high-resolution tissue-wide cell type map. To construct regional ground truth rankings, we applied an 8 × 8 sliding window across the tissue, mirroring our model’s inference approach. For each windowed region, we aggregated the cell type classifications of all enclosed spots. If a region contained more than 10 valid spots, we computed the Shannon entropy of the spot classification distribution—weighted by log(1 + *n*_spots_)—to quantify cell type complexity. For tumor-targeted evaluation, we scored each region based on the total number of tumor-classified spots.

We evaluated our model’s ROI rankings against these ground truth measures using three standard metrics:

- **Spearman’s rank correlation (***ρ*): Measures global concordance between model and ground truth rankings, indicating overall prioritization accuracy^51^.
- **Top-K overlap (Overlap@K)**: Computes the percentage overlap between the top K regions identified by the model and by the ground truth, highlighting performance at the high-priority end of the list.
- **Intersection over Union (IoU@K)**: Assesses spatial alignment by measuring the area-based overlap between the top K predicted and ground truth regions^52^.

## 3. Results

We assessed our hybrid model on four distinct Visium HD tissue types, evaluating both cell diversity and tumor presence. Across the board, our model consistently outperformed baseline methods in identifying biologically meaningful regions of interest.

### 3.1. SpatialFinder Hybrid Model Surpasses VLM Baselines

To establish a quantitative reference point, we benchmarked SpatialFinder against three prominent vision-language models: PLIP and CONCH^53^—state-of-the-art VLMs in pathology—and MUSK, the vision-language component of our own hybrid system.

All models were evaluated using an identical preprocessing and inference pipeline, including tissue cropping, 224 × 224 patching, background filtering, UMAP-based dimensionality reduction, and an 8 × 8 sliding window for scoring regions of interest (ROIs). The critical distinction is that baseline models do not incorporate human-in-the-loop cellular insights: they exclude nuclei segmentation and pathologist-informed cell classification. For the diversity task, baseline scoring relied solely on the UMAP spread of patch embeddings (*w*=1), while the targeted task used only text-to-image similarity (*w*=1), computed via the 90th-percentile cosine similarity between patch embeddings and the text prompt “tumor.”

PLIP and CONCH are used strictly zero-shot as plug-in encoders, while MUSK (VLM-only) serves as an internal ablation to isolate the added value of the classifier. Though not shown, random ROI selection performs substantially worse than all tested methods and is omitted for clarity; our aim is to benchmark against strong, modern VLM-only pipelines and demonstrate the performance gains from integrating expert-driven cellular information.

As detailed in **Table 1**, SpatialFinder matched or outperformed baselines across all tissue types for both diversity and tumor detection tasks^c^. In the colon dataset, it achieved a Spearman’s *ρ* of 0.89, and showed nearly 80% overlap with ground truth in the top 10% of ranked regions in the prostate set.

**Table 1:** Performance of SpatialFinder against other baseline VLMs across four datasets. Bold texts indicate best performed results.

|  | Model | Cell Diversity |  |  | Tumor |  |  |
| --- | --- | --- | --- | --- | --- | --- | --- |
| | | $\rho$ | Overlap@10% | IoU@10% | $\rho$ | Overlap@10% | IoU@10% |
| Prostate | PLIP | 0.484 | 2.92% | 0.169 | 0.222 | 23.49% | 0.278 |
|  | CONCH | 0.434 | 1.23% | 0.147 | 0.829 | 44.15% | 0.486 |
|  | MUSK | 0.650 | 38.68% | 0.478 | 0.833 | 53.87% | 0.553 |
|  | SpatialFinder | **0.762** | **47.45%** | **0.559** | **0.857** | **78.77%** | **0.775** |
| Lung | PLIP | 0.458 | 8.82% | 0.249 | 0.254 | 0.00% | 0.062 |
|  | CONCH | 0.521 | 12.19% | 0.269 | 0.826 | 35.08% | **0.492** |
|  | MUSK | 0.594 | 17.24% | 0.366 | 0.653 | 1.98% | 0.179 |
|  | SpatialFinder | **0.653** | **25.27%** | **0.406** | **0.886** | **37.26%** | 0.401 |
| Colon | PLIP | 0.167 | 13.20% | 0.172 | 0.352 | 11.30% | 0.217 |
|  | CONCH | 0.161 | 23.70% | 0.249 | 0.619 | 15.70% | 0.309 |
|  | MUSK | 0.196 | 31.90% | 0.324 | 0.527 | 11.70% | 0.243 |
|  | SpatialFinder | **0.365** | **38.10%** | **0.445** | **0.623** | **19.10%** | **0.355** |
| Kidney | PLIP | 0.660 | 32.26% | 0.445 | — | — | — |
|  | CONCH | 0.639 | 33.77% | 0.482 | — | — | — |
|  | MUSK | 0.701 | 37.36% | 0.540 | — | — | — |
|  | SpatialFinder | **0.801** | **48.30%** | **0.594** | — | — | — |

SpatialFinder’s robust performance across lung, colon, prostate, and kidney datasets underscores its versatility. While CONCH slightly outperformed it in IoU on the lung tumor benchmark—highlighting that VLM-only approaches can excel in some targeted tasks—our hybrid approach consistently delivered stronger results overall. These findings affirm the benefit of fusing semantic visual features with granular cellular context, offering a more reliable and interpretable alternative to vision-only methods.

### 3.2. Consistent Performance Across Top-K Regions

To evaluate the model’s robustness in identifying high-priority regions, we analyzed its Top-K performance by varying the value of K. **Figure 5** illustrates the Top-K Overlap scores for our model and the baselines, with K varying from 10 to 1000.

**Fig. 1:**
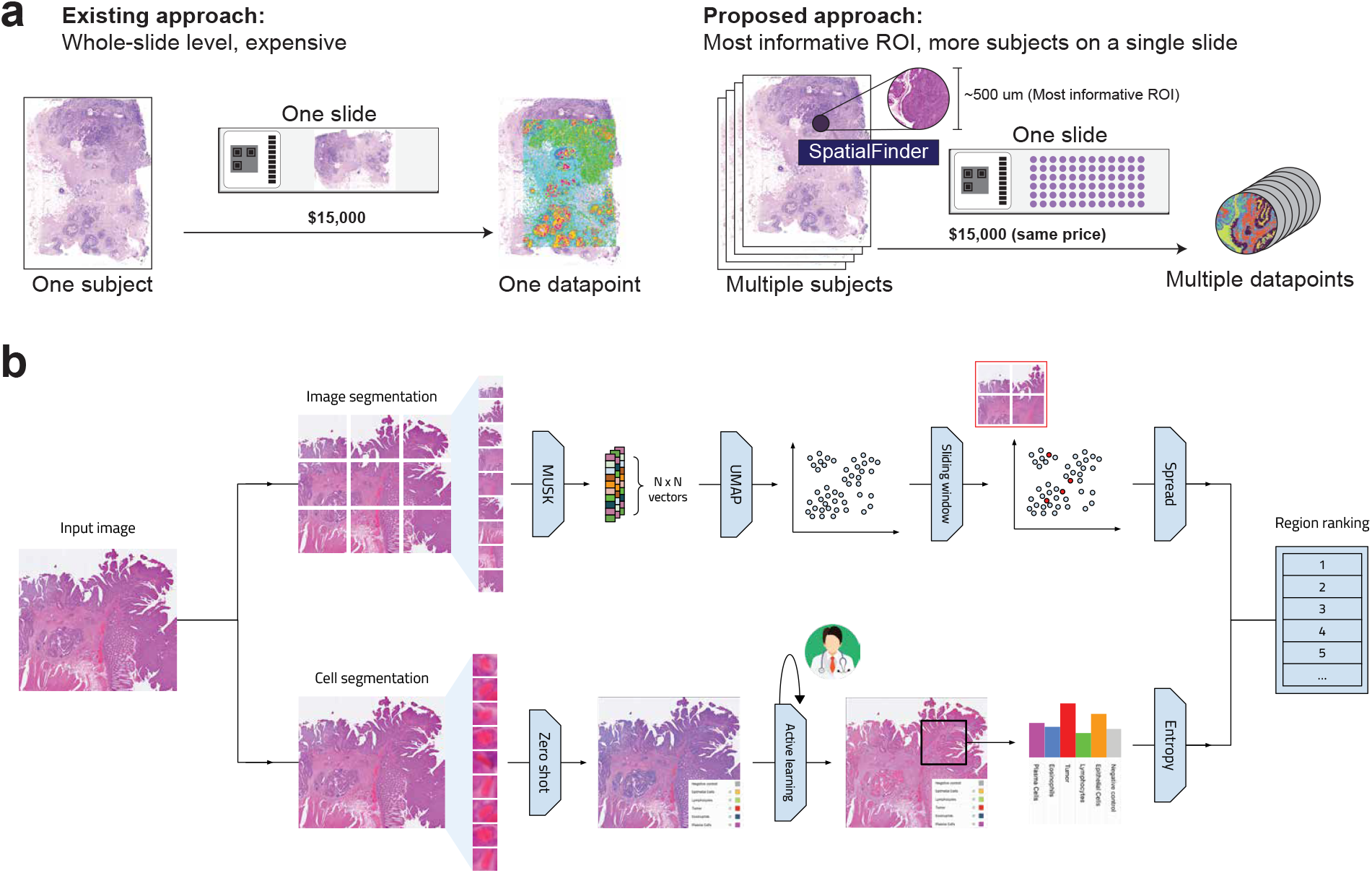
(a) Current spatial omics cost upwards of $15,000 per slide, creating a high cost bottleneck for broader application. Our proposed alternative addresses this by identifying only the most relevant regions within each slide for gene sequencing, dramatically increasing the number of subjects that can be analyzed at the same cost. (b) An overview of our approach. SpatialFinder utilizes a sliding-window procedure on the input image to extract adjacent patches for analysis. Image patches are encoded into high-dimensional feature vectors, which are then projected into a lower-dimensional space via UMAP for visualization and evaluation. In parallel, we train a nuclei classifier through an active-learning loop with expert pathologist input to classify cells accurately from H&E. Regions are ranked by the weighted scores from both arms of the proposed pipeline.

**Fig. 2:**
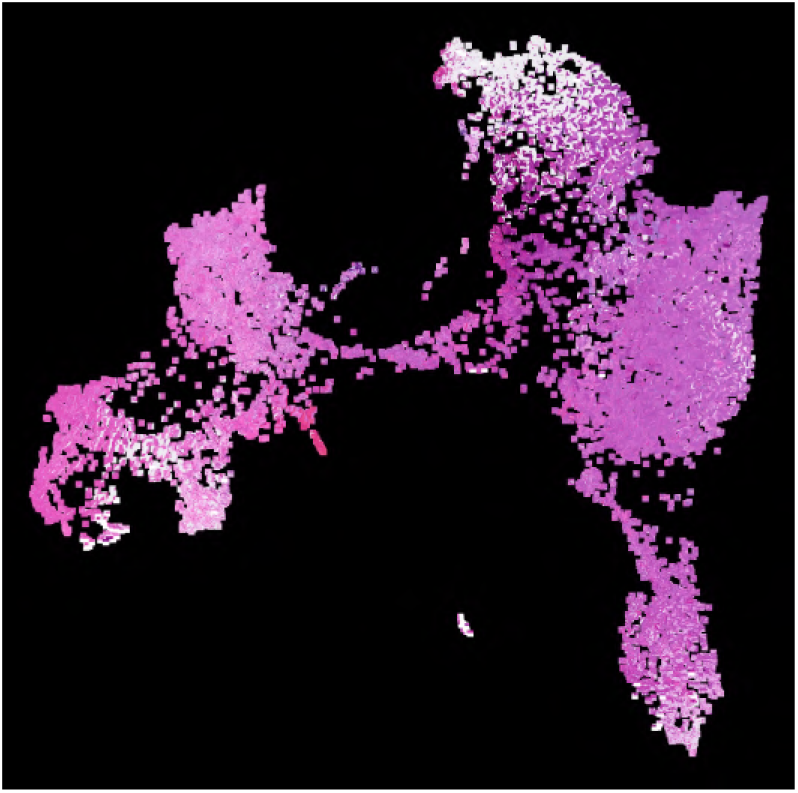
UMAP visualization in 2 dimensions of a tissue slide’s patch embeddings with the patch images superimposed.

**Fig. 3:**
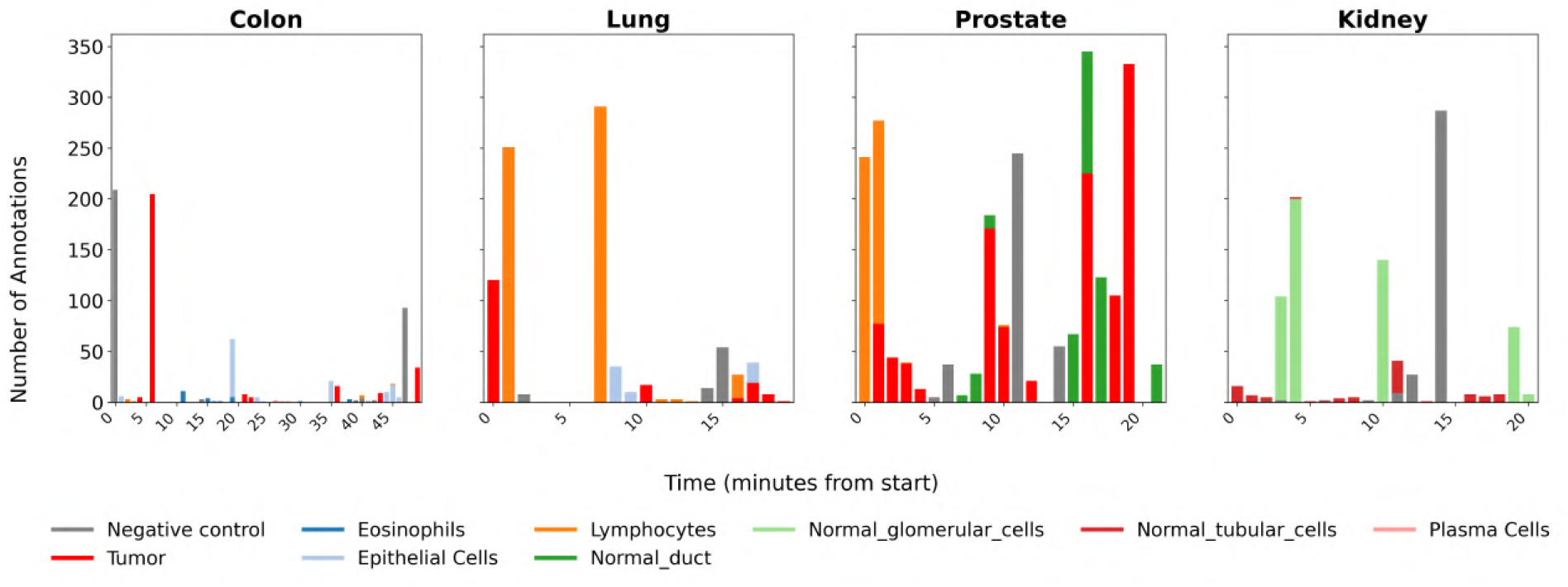
Annotation timestamps for the 4 Visium HD tissue types used to train our active learning classifiers. Annotation sessions lasted for an average of 27 minutes.

**Fig. 4:**
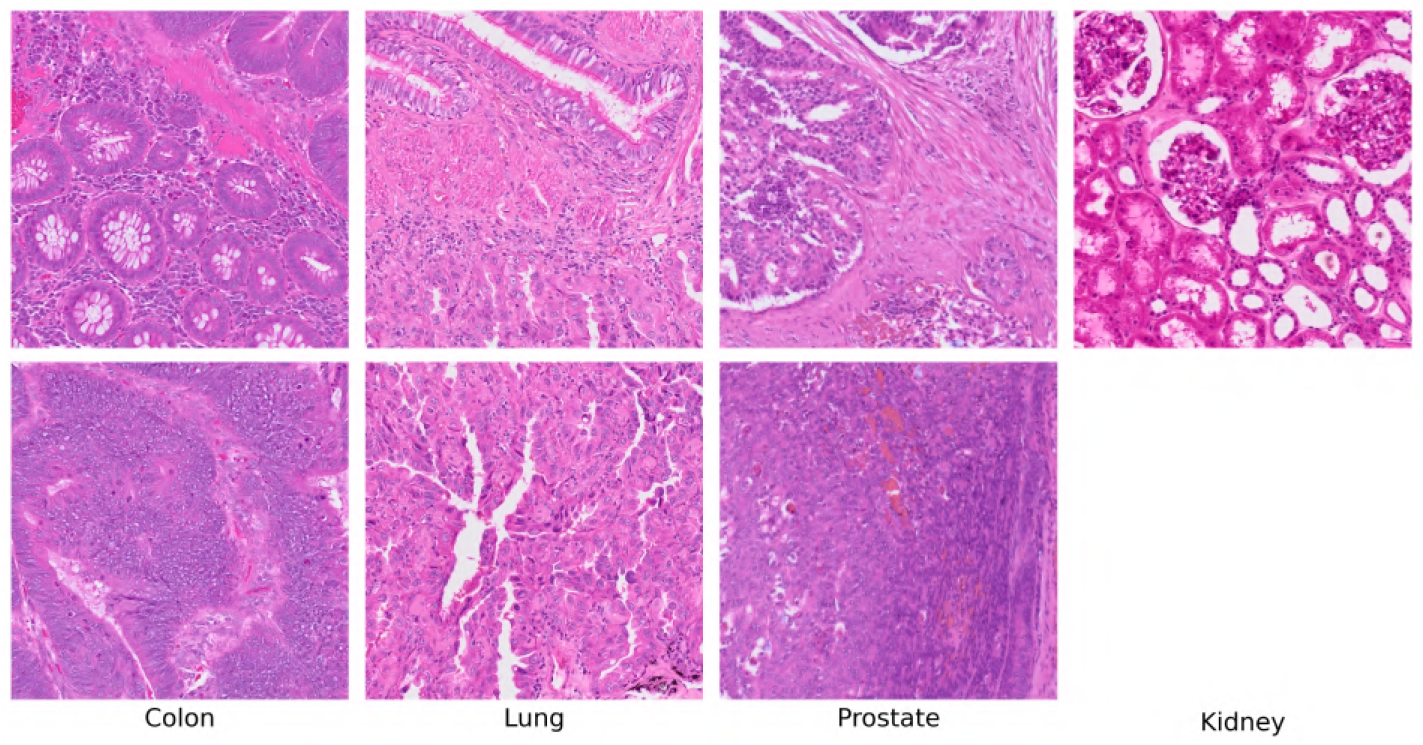
Row 1 displays the top results for cellular diversity. Row 2 is conditioned on tumor presence. The kidney sample is non-cancerous and therefore does not have a tumor entry.

**Fig. 5:**
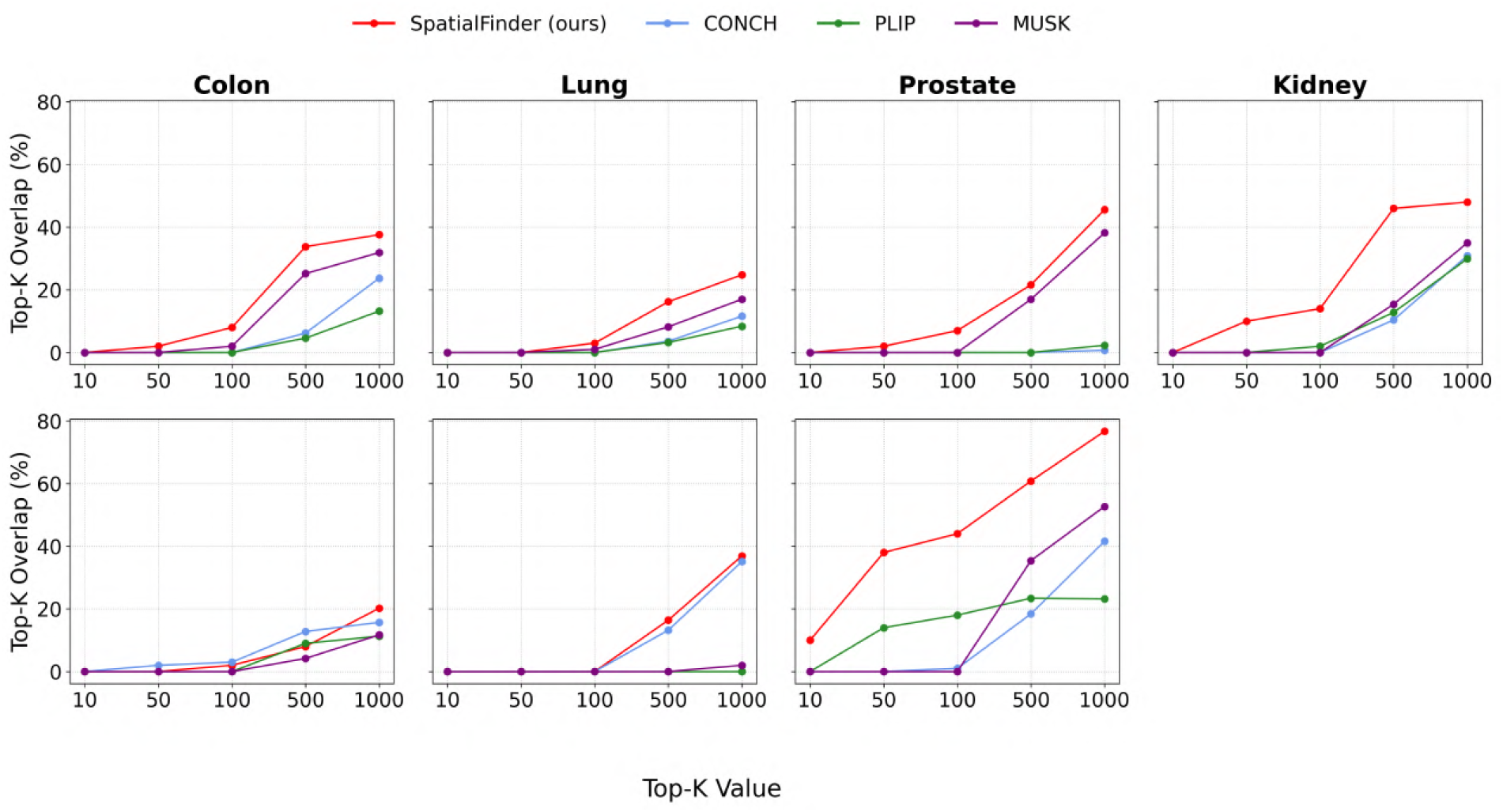
Top-K performance on cell diversity benchmark varied over multiple K. SpatialFinder consistently achieves higher selection accuracy. Top row: cell diversity. Bottom row: tumor presence.

Our model maintains superior performance throughout the entire range, largely surpassing the baselines at both large-scale evaluations (K=*{*500, 1000*}*) and at more focused, decision-critical levels (K=*{*50, 100*}*). Notably, in the prostate tissue, it even accurately identifies a subset of the Top-10 ground truth regions. These results highlight the model’s ability to reliably identify a compact, high-confidence set of candidate regions out of about 10,000 options, offering practical advantages for targeted downstream analyses.

### 3.3 Qualitative Agreement with Ground Truth Spatial Patterns

In addition to quantitative metrics, we qualitatively assessed our model’s ability to capture biologically meaningful spatial patterns. **Figure 6** compares heatmaps of score predictions against ground truth distributions for the prostate tissue.

**Fig. 6:**
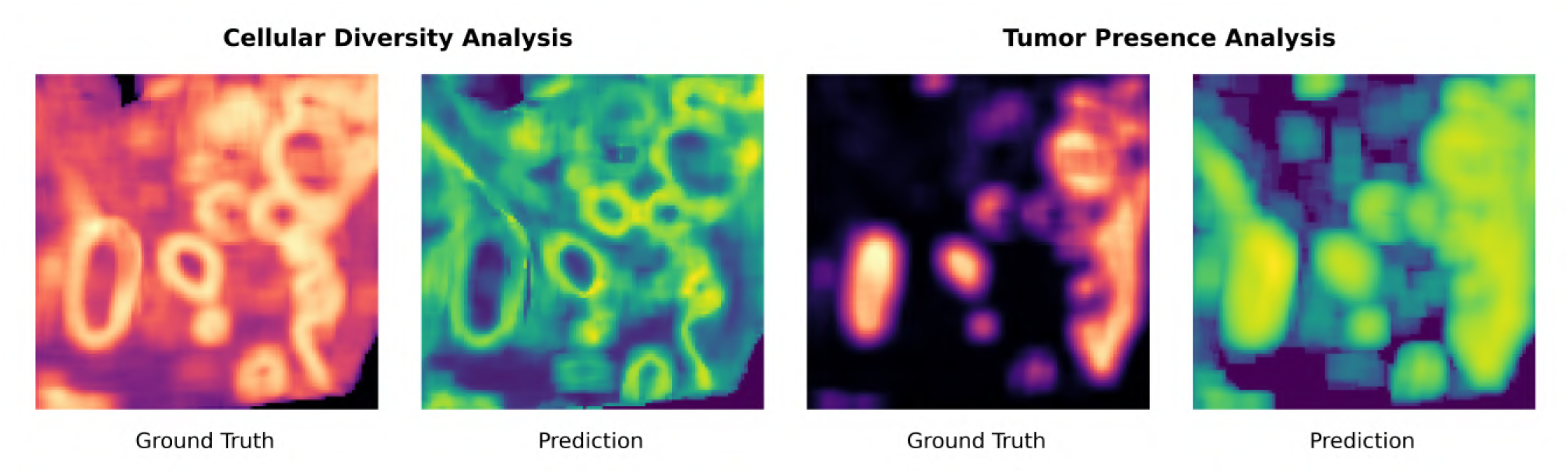
Comparison of predicted versus ground truth heatmaps for cell diversity (left) and tumor presence (right). Each pixel corresponds to the centroid of an 8 × 8 patch, colored according to the associated score. Brighter pixels represent higher scores. Ground truth is derived from actual ST data; prediction is based on H&E data only.

The visual alignment is compelling: predicted tumor hotspots closely match high-scoring ground truth regions. The model also reliably identifies heterogeneous tissue areas, capturing the complex cellular architecture underlying transcriptomic diversity. Notably, it highlights regions at the boundary of tumor-dense zones, areas known to serve as key sites of tumor-immune interaction^54^. Pinpointing this boundary is a central motivation in spatial transcriptomics, as these areas often represent critical sites of immune infiltration, tumor evolution, and therapeutic response^55^. The model’s attention towards these biologically relevant regions underscores its ability to uncover meaningful spatial patterns beyond simple statistical correlations.

In our experiments, we evaluated for cell diversity and tumor presence separately to assess their individual performance. However, they can also be combined or conditioned on one another to support more nuanced or multi-objective selection criteria. For instance, we also explored a two-step approach where we first applied a gating threshold on the Targeted Conditional Score to isolate tumor-enriched regions, and then ranked those regions by their Regional Diversity Score to prioritize the most heterogeneous tumor microenvironments. **Figure 7** illustrates this with a heatmap of the predicted cell diversity, where each pixel represents an 8 × 8 patch square and its color quantifies the diversity score.

**Fig. 7:**
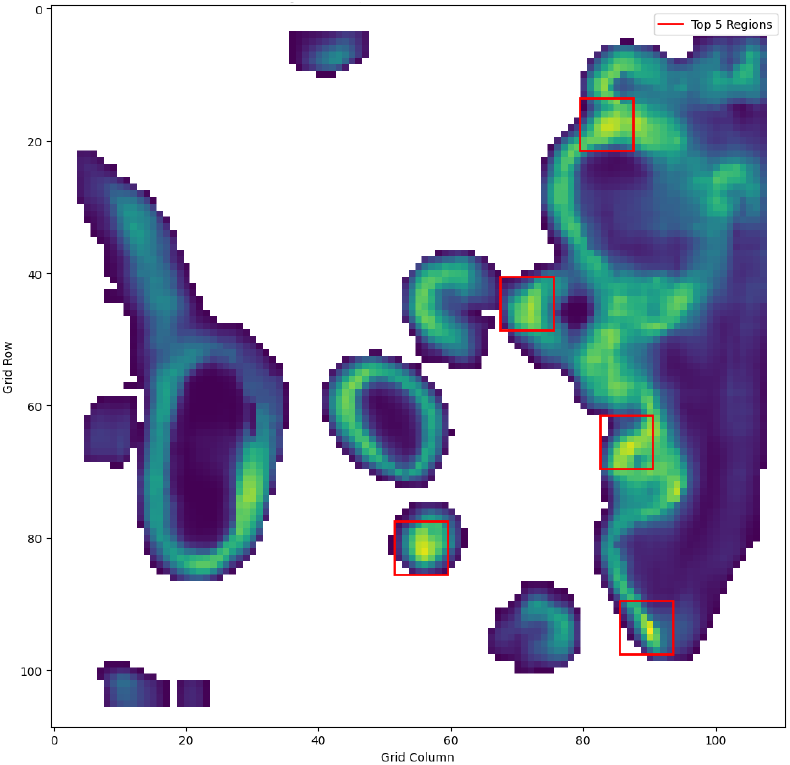
A heatmap of our integrated model with the tumor threshold.

### 3.4. Synergy of Cellular and Visual Features Drives Performance

A central premise of our approach is that integrating cellular and visual-semantic information leads to better performance than either source alone. **Figure 8** illustrates this effect by plotting Spearman’s *ρ* as we vary the weighting parameter from 0.0 (classifier-only) to 1.0 (VLM-only). In the majority of these cases, the highest performance is achieved with an intermediate weighting, demonstrating a meaningful synergy between modalities. Even away from these optima, the blended model largely stays above other fixed-weight baselines, underscoring its robustness to imperfect weighting.

**Fig. 8:**
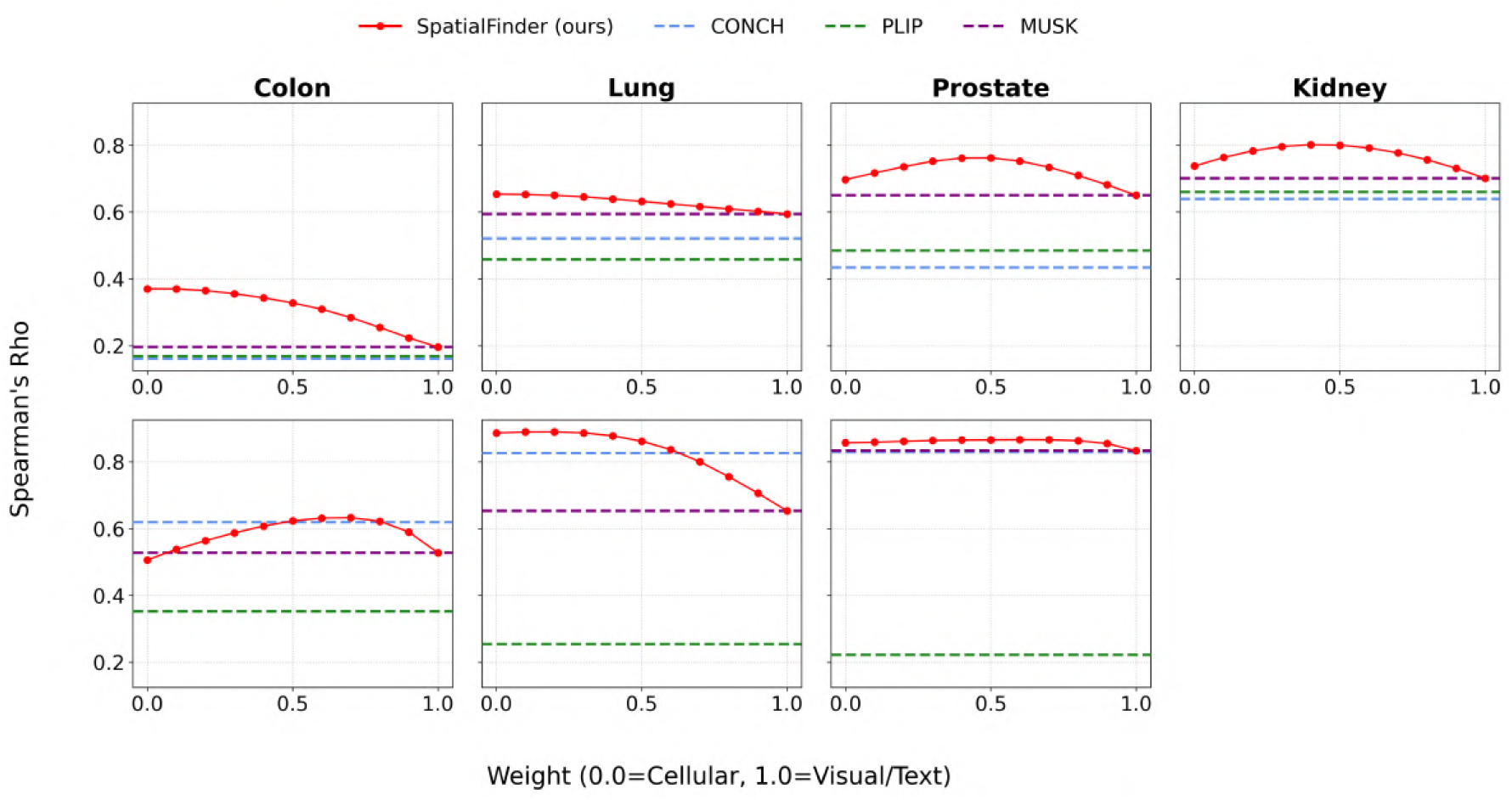
Spearman’s *ρ* as a function of the weighting parameter between classifier and VLM outputs. Top row: cell diversity. Bottom row: tumor presence.

This complementary effect is further illustrated in **Figure 9**, which shows how Top-K IoU varies with the weighting parameter. Across multiple values of K, the best performance consistently arises from a hybrid representation, reinforcing the idea that cellular and visual features offer distinct and synergistic contributions. At lower values of K, the cell classifier tends to perform better, indicating its strength in pinpointing the most salient regions.

**Fig. 9:**
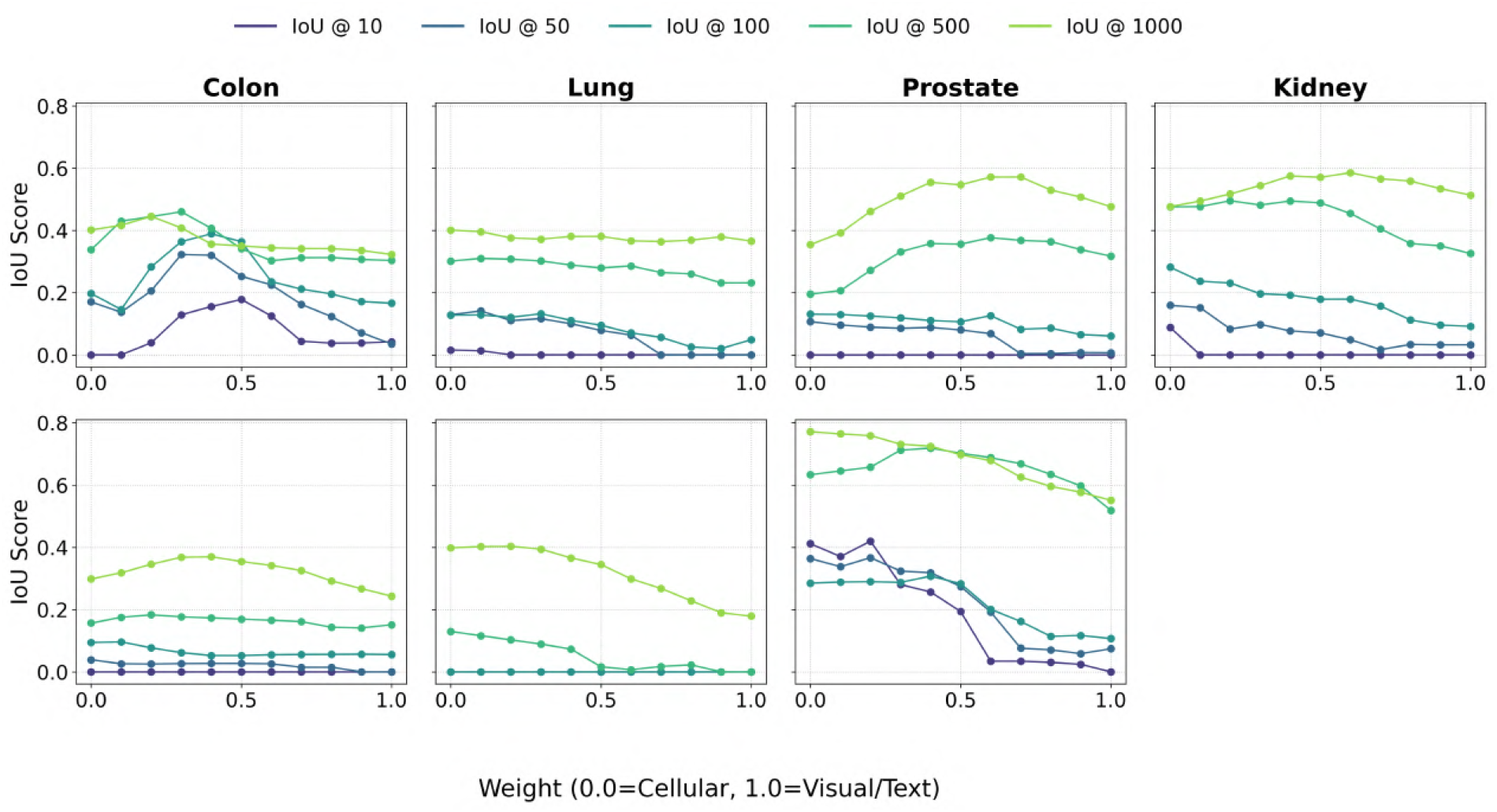
Intersection over Union scores for multiple K-values across varying weights for our model. Top row: cell diversity. Bottom row: tumor presence.

### 3.5. Pathologist-Guided Tuning Rapidly Boosts Performance

To evaluate the efficiency of human-in-the-loop refinement, we conducted an ablation study measuring model performance as a function of annotation time. Starting from a zero-shot classifier, we incrementally reintroduced pathologist-labeled data in timed intervals. **Figure 10** displays the resulting performance trajectory on both cell diversity and tumor benchmarks.

**Fig. 10:**
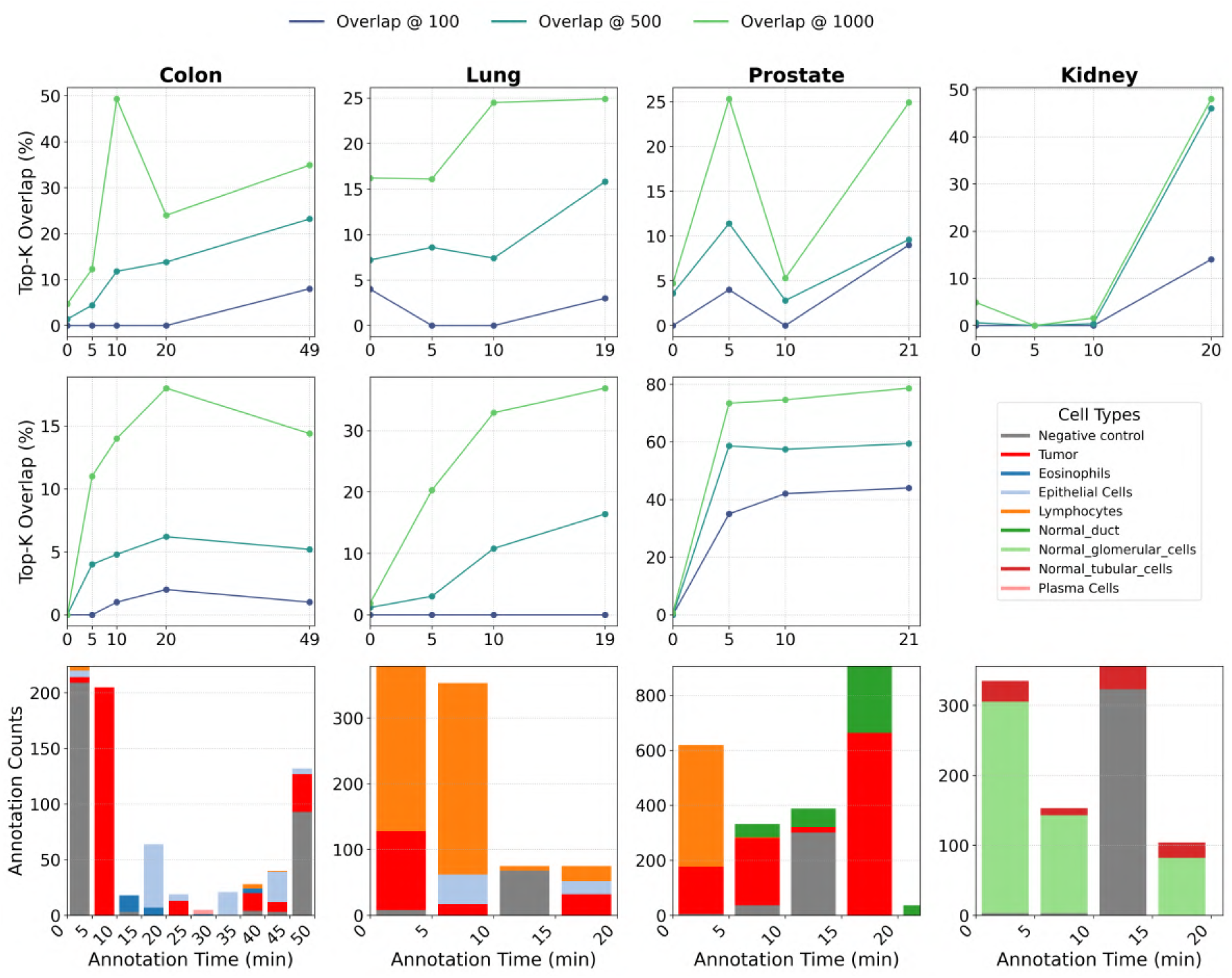
SpatialFinder’s performance on Top-K as pathologist annotation time increases. Top row: cell diversity. Middle row: tumor presence. Bottom row: annotation counts over time.

The results show that while the zero-shot model provides a strong initialization, even minimal expert input significantly improves performance. In most tissue, as little as 5-10 minutes of annotation captures the majority of the total gain, particularly for broader retrieval tasks such as Top-500 or Top-1000 overlap. For example, in colon and prostate tissue, short annotations recover over 50% of the Top-1000 list, whereas kidney requires more sustained annotations (10-20 minutes) before a sharp performance boost is observed.

Across all benchmarks, our framework consistently outperforms the zero-shot baseline. These findings underscore its practical value: with just a few minutes of expert annotation, we can train a high-performing, tissue-specific model, enabling fast, expert-guided adaptation in real-world clinical and research environments.

## 4. Conclusion and Future Directions

We introduced SpatialFinder, a framework that combines vision-language models with pathologist-guided cell classification to identify informative subregions for spatial transcriptomics. By integrating high-level semantic features from a VLM with fine-grained cellular labels derived through active learning, our method enables targeted selection of small 500×500μm regions likely to capture transcriptomic heterogeneity. We demonstrate its flexibility across both discovery-driven (diversity) and hypothesis-driven (targeted) use cases, offering a practical strategy for reducing sequencing costs while preserving biological insight.

### 4.1. Limitations

A core challenge in this domain is defining what constitutes an “interesting” region^56^. Different use cases (e.g. tumor profiling vs. immune mapping) may demand different criteria. We addressed this by exploring both cell diversity and tumor enrichment, but these proxies remain limited. In particular, measuring diversity via our proposed methods may miss subtler or more specific compositional signals. Future work could incorporate deconvolution tools such as cell2location^57^ or RCTD^58^ to better resolve cellular heterogeneity.

Our evaluation relies on ranking metrics like Spearman’s *ρ* and Overlap@K, which are effective for global assessment but may not reflect performance in top-few selection scenarios. There is a tradeoff between optimizing overall ranking and prioritizing the highest-scoring ROIs, which could be better addressed with retrieval-style, top-K-oriented objectives that are now emphasized in visual–omics benchmarks^59,60^. Robustness in such settings also depends on prompt design: instance-specific prompting, as in QAP, has been shown to outperform task-agnostic or pooled approaches by tailoring inputs to tissue-derived features^61^.

### 4.2. Future Work

Looking ahead, scaling pathology-specific vision–language models may further improve region selection in spatial transcriptomics. Models like PLIP and CONCH already show strong performance in zero-shot tissue localization tasks^39,53^, suggesting their potential for identifying biologically informative areas without retraining. Incorporating protein-level validation (e.g., imaging mass cytometry) may help confirm the molecular relevance of predicted regions^62^. As sequencing technologies vary widely in resolution and format, developing shared benchmarks will be important for comparing model outputs across platforms^63^. In addition, iterative sequencing could make ROI selection adaptive, allowing models to refine their predictions based on early gene expression feedback^64,65^.

Finally, we envision adding a third reasoning layer via large language models (LLMs). Our framework leverages VLMs and classifier-driven scores; adding a generative LLM component could enable natural language queries like “tumor borders infiltrated by immune cells,” and allow the model to interpret, rank, and describe these ROIs back to the user. Pathology-specific multimodal assistants already show that coupling a vision encoder with a large language model enables natural-language question answering over slides and interactive assistance^66^.

In summary, SpatialFinder offers a step towards scalable, intelligent spatial biology. By combining VLMs with expert-in-the-loop classification, it opens a flexible, extensible foundation for guiding spatial transcriptomic experiments more efficiently and meaningfully.

## Supporting information

Figure 1: Overview of our approach

## 5. Acknowledgments

We thank Yinuo Xu for help with cell type labeling, Tiancheng Bao for data preprocessing support, and Flaticon for the doctor clipart in Figure 1.

## Footnotes

a While the Visium HD platform also offers a higher 2 × 2μm resolution, we found this to be sub-cellular and thus susceptible to gene expression spillover between adjacent cells.

b A full list of marker genes is available at: https://gist.github.com/jonrxu/63e9319016c257a54c823fbaab26bb27

c The Visium HD kidney sample is non-cancerous and therefore excluded from tumor evaluations.

