## Supplementary figures and images for "SpatialFinder: A Human-in-the-Loop Vision-Language Framework for Prioritizing High-Value Regions in Spatial Transcriptomics"

### Figure 1: Overview of our approach

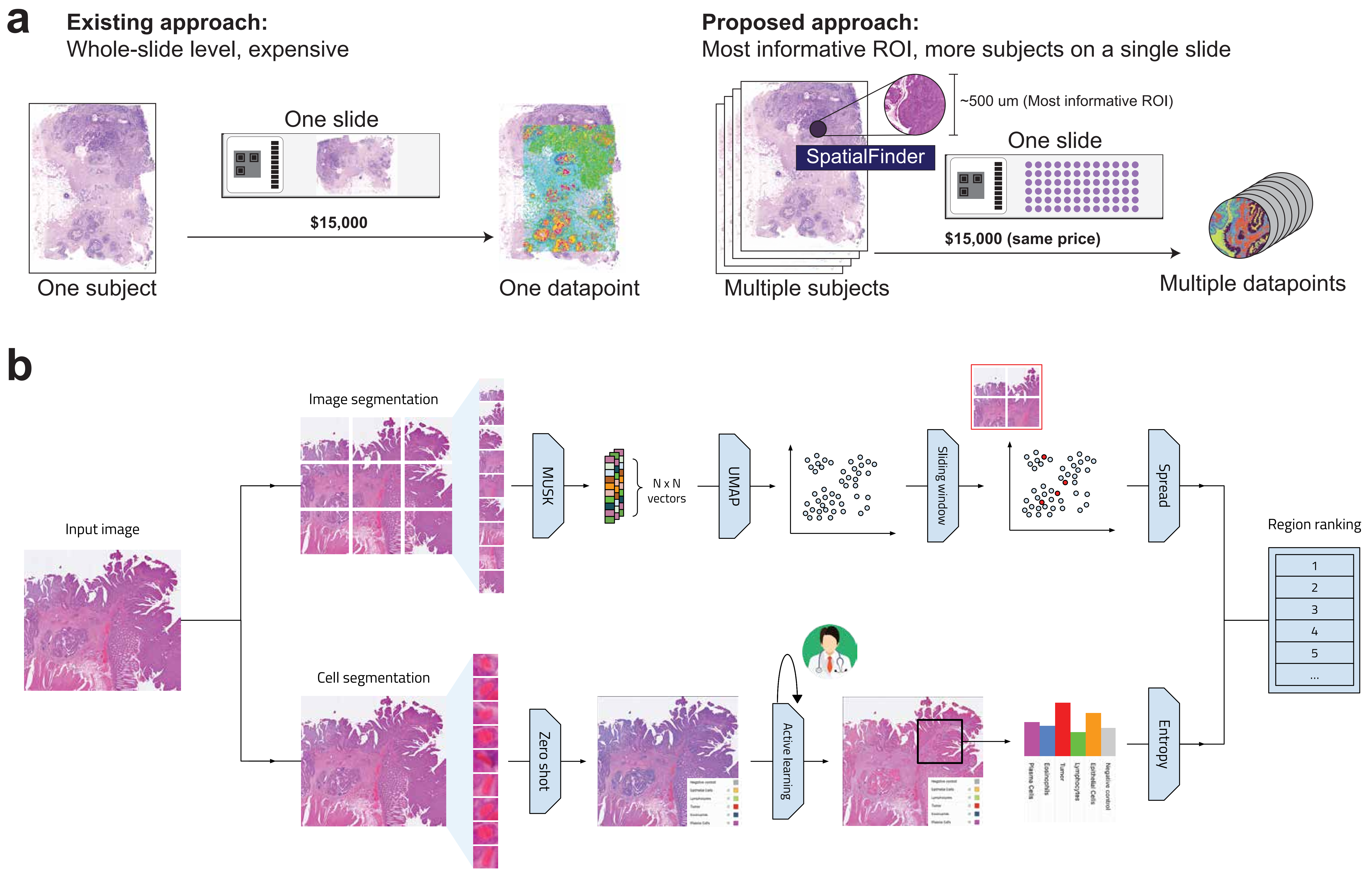
